# The regulatory control of *Cebpa* enhancers and silencers in the myeloid and red-blood cell lineages

**DOI:** 10.1101/369553

**Authors:** Andrea Repele, Shawn Krueger, Michelle Y. Tuineau, Manu

## Abstract

During development, cell identity is determined by lineage-specific transcriptional programs encoded in the *cis*-regulatory DNA sequence of developmental genes. The sequence-level regulatory logic—identities of bound transcription factors (TFs), TF binding sites, and TF regulatory roles—of most developmental *cis*-regulatory modules (CRMs) is yet to be determined. We had previously developed an approach for inferring regulatory logic *de novo* by training sequence-based thermodynamic models on comprehensive reporter activity and gene expression datasets and applied it to *Cebpa*, an important hematopoietic gene. Here, we experimentally test thermodynamic models to decode the *cis*-regulatory logic of 4 enhancers and 3 silencers neighboring *Cebpa* at the resolution of individual binding sites. *Cebpa* is expressed at high and intermediate levels in neutrophils and macrophages respectively and downregulated in non-myeloid lineages. We tested the binding sites and functional roles of inferred TFs by designing and constructing mutated CRMs and comparing theoretical predictions of their activity against empirical measurements. Reporter activity was measured in PUER cells, which can be induced to differentiate into macrophages or neutrophils. All four enhancers were found to be simultaneously active in undifferentiated PUER cells and early-stage macrophages and neutrophils, and activated by combinations of PU.1, C/EBP family TFs, Egr1, and Gfi1. We show that silencers repress the activity of the proximal promoter in a dominant manner in G1ME cells, which are derived from the red-blood cell lineage. Dominant repression in G1ME cells can be traced to binding sites for GATA and Myb, a motif shared by all of the silencers. Finally, we demonstrate that GATA and Myb act redundantly to silence the proximal promoter. The result that silencers quench the promoter selectively in non-myeloid cells indicates that dominant repression is a novel mechanism for resolving hematopoietic lineages. Furthermore, *Cebpa* has a fail-safe *cis*-regulatory architecture, featuring several functionally similar CRMs, each of which contains redundant binding sites for multiple TFs. Lastly, by experimentally demonstrating the predictive ability of our sequence-based thermodynamic models, this work highlights the utility of this computational approach for decoding the logic of mammalian gene regulation.

## Introduction

Cell-fate decisions during hematopoiesis are dictated by gene regulatory networks (GRNs Singh *et al*., 2014; Laslo *et al*., 2008; Enver *et al*., 2009) that switch state based on the regulatory activities of transcription factors (TFs) and cytokine signals (Rieger *et al*., 2009; Mossadegh-Keller *et al*., 2013). Hematopoietic lineage resolution has often been interpreted in the context of a simple network motif, the bistable switch (Huang *et al*., 2007; Enver *et al*., 2009; Laslo *et al*., 2006). Functional genomics data however suggest a more complex, densely interconnected, organization of hematopoietic GRNs. First, network reconstructions based on genome-wide gene expression data reveal large modules of co-regulated genes (Novershtern *et al*., 2011). Second, genome-wide TF binding data show that most loci, and often individual sites, are co-bound by multiple TFs (Wilson *et al*., 2010; Nègre *et al*., 2011). If hematopoietic GRNs were indeed densely interconnected, it would imply that the bistable switch framework is inadequate for modeling and predicting cell-fate choice. Although co-expression or co-binding data are highly suggestive of densely connected GRNs, the interactions must be established at the *cis*-regulatory level for delineating the networks with high confidence. With the exception of a few genes that have been well-characterized in this regard (Wilson *et al*., 2011), the regulatory logic of most hematopoietic genes remains unknown. In this study, we decode the regulatory logic of seven *cis*-regulatory elements (CRMs) of *CCAAT/Enhancer binding protein,α* (*Cebpa*) at binding-site resolution aduring myeloid differentiation.

*Cebpa* encodes a TF that is necessary for neutrophil development (Zhang *et al*., 1997) as well as the specification of hepatocytes and adipocytes (Legraverend *et al*., 1993; Rosen *et al*., 2002). During hematopoiesis, *Cebpa* is expressed in hematopoietic stem cells, granulocyte-monocyte progenitors (GMPs), neutrophils, and macrophages (http://biogps.org/gene/12606; Wu *et al*., 2016; Lattin *et al*., 2008). Although the most apparent hematopoietic phenotype of *Cebpa*^*-/-*^mice is neutropenia (Zhang *et al*., 1997), *Cebpa* also has a role in specifying macrophages. *Cebpa* is expressed at intermediate and high levels in macrophages and neutrophils respectively and the cell-fate decision is thought to depend on the ratio of PU.1, a TF necessary for white-blood cell lineages (Scott *et al*., 1994), and C/EBP*α* expression levels (Dahl *et al*., 2003). Correspondingly, the cell-fate decision has been modeled as a bistable switch in which PU.1 and C/EBP *α* activate the mutual antagonists *Egr1/2* and *Gfi1* respectively (Laslo *et al*., 2006). *Cebpa* is also sufficient for specifying macrophages, since B-cells can be transdifferentiated into them by expressing *Cebpa* ectopically (Xie *et al*., 2004).

Despite its essential and pleiotropic functions, the *cis* regulation of *Cebpa* during hematopoiesis is poorly understood. C/EBP*α*, C/EBP*β*, and C/EBP*δ* are known to activate *Cebpa* by binding to its proximal promoter (Legraverend *et al*., 1993). Recently, ZNF143 was shown to bind and activate the promoter in a human myeloid cell line (Gonzalez *et al*., 2017). *Cebpa* is regulated in 32Dcl3 myeloid cells by PU.1, other Ets TFs, SCL, Gata2, Myb, and C/EBP*α*, which bind to an enhancer located 37kb downstream of the gene (Cooper *et al*., 2015; Guo *et al*., 2016). It is not known whether, like other pleiotropic TFs (Wilson *et al*., 2011), *Cebpa* is also regulated by multiple CRMs. More importantly, it is not understood how the regulatory contributions of these and other TFs modulate *Cebpa*’s gene expression during differentiation.

The deficits in our understanding of *Cebpa*’s regulatory logic illustrate the general challenge of decoding gene regulation on a genomic scale. The challenge arises from the complexity of gene regulation—genes may be regulated by multiple CRMs (Wilson *et al*., 2011; Spitz & Furlong, 2012) and each CRM may, in turn, be jointly regulated by several TFs exerting positive or negative influence over the target gene (Small *et al*., 1992; Janssens *et al*., 2006; Heinz *et al*., 2010; ENCODE Project Consortium *et al*., 2012; Gerstein *et al*., 2012, 2010). The problem of decoding regulatory logic, therefore, is one of mapping multiple inputs—TF concentrations—to a single output—the rate of transcription.

We have developed a computational approach to solve the problem of mapping multiple TF inputs to transcriptional output and decoding regulatory logic (Bertolino *et al*., 2016). We utilize sequence-based models of transcription (Janssens *et al*., 2006; Segal *et al*., 2008; Kim *et al*., 2013) that simulate gene regulation by multiple TFs according to precise mechanistic rules of TF-DNA binding, competition, repression, and cooperation (Arnosti *et al*., 1996a; Cantor & Orkin, 2001, 2002; He *et al*., 2012; Heinz *et al*., 2010; Kulkarni & Arnosti, 2005; Small *et al*., 1993, 1996). The model take estimates of TF concentrations, CRM DNA sequence, and position weight matrices as inputs and computes the resulting CRM activity as an output. Our approach does not require *a priori* knowledge of the identities or the regulatory roles, activation or repression, of the TFs regulating a CRM. The TFs regulating a CRM and their regulatory roles are inferred *in silico* by testing many alternative models, each realizing a potential regulatory scheme, against quantitative reporter data. The composition of the best fitting model then implies the regulatory roles of the TFs most congruent with the observed patterns of CRM-and cell-type-specific reporter activity. It is worth noting that this procedure not only produces a description of the TFs, their roles, and their binding sites, but also yields predictive models of CRM function.

We previously applied our approach to *Cebpa* in order to comprehensively decode its regulation (Bertolino *et al*., 2016) during macrophage-neutrophil differentiation. The reporter assays were carried out in PUER cells (Walsh *et al*., 2002), which act as bipotential granulocyte-monocyte progenitors (GMPs) and can be differentiated into macrophages or neutrophils by treatment with 4-OH-tamoxifen (OHT) in the presence of IL-3 or G-CSF respectively (Dahl *et al*., 2003; Laslo *et al*., 2006). We identified 8 CRMs, of which 7 are novel, lying between −39kb and +38kb from the *Cebpa* transcription start site (TSS). Four CRMs, including one encompassing the +37kb enhancer identified by Guo *et al.* (2012), acted as enhancers and upregulated reporter activity 2-to 6-fold relative to the proximal promoter of *Cebpa*. The remaining CRMs, appeared to behave as silencers and repressed the activity of the proximal *Cebpa* promoter in a dominant fashion.

Our computational analysis inferred a comprehensive map of the regulation of the *Cebpa* locus and suggested a novel mechanism of lineage resolution (Bertolino *et al*., 2016). The enhancers were predicted to be activated by PU.1, C/EBP family TFs, Egr1, and Gfi1 and repressed by Myb. Surprisingly, the model predicted that the silencers exert repression through the activity of TFs strongly expressed in non-myeloid cell types, GATAs, Ebf1, and Myb. The silencing of *Cebpa* is necessary for the specification and maintenance of non-myeloid cell fates (Chou *et al*., 2009; Heinz *et al*., 2010; Xie *et al*., 2004). Dominant repression of the *Cebpa* promoter by distal silencers in non-myeloid lineages therefore might be a mechanism for resolving lineages. These inferences must however be regarded as predictions since they are yet to be verified experimentally.

Here, we report the regulatory logic of 4 enhancers and 3 silencers of *Cebpa* at binding-site resolution. We determined the regulatory logic of these CRMs by rigorously testing the predictions of our computational models. We predicted the effect of mutations to one or more binding sites by simulating the regulation of mutated DNA in the model. The predictions were experimentally tested by synthesizing mutated CRMs and assaying their activity. The regulatory logic of enhancers was investigated in PUER cells, representing the myeloid lineage where *Cebpa* is expressed robustly. The function and regulation of silencers was investigated in G1ME cells (Stachura *et al*., 2006), which are derived from *Gata1*^*-/-*^ mice and represent the red-blood cell lineage by virtue of being blocked at the megakaryocyte-erythrocyte progenitor (MEP) stage.

Using this combination of computational and experimental regulatory analysis we show that *Cebpa* has a complex and redundant *cis*-regulatory architecture. All four enhancers are simultaneously active in undifferentiated PUER cells and early-stage macrophages and neutrophils. Using mutagenesis, we demonstrate that PU.1 activates CRM *Cebpa(16)*, C/EBP family TFs activate CRMs *Cebpa(7)* and *Cebpa(16)*, while Gfi1 and Egr1 activate CRMs *Cebpa(7)* and *Cebpa(14)* respectively. We show that silencers repress the activity of the proximal promoter 3-fold in a dominant manner in G1ME cells but are neutral in PUER cells. Dominant repression in G1ME cells can be traced to GATA and Myb sites, a common motif present in all three of the validated silencers. Finally, we demonstrate through a detailed regulatory analysis of one of the silencers, *Cebpa(11)*, that GATA and Myb act redundantly to silence the proximal promoter.

## Materials and Methods

### Cell culture

We utilized *Spi1*^*-/-*^ cells, expressing conditionally activable PU.1 protein, which can be differentiated into macrophages or neutrophils by PU.1 activation (PUER Dahl *et al*., 2003; Laslo *et al*., 2006; Walsh *et al*., 2002). PUER cells were routinely maintained in complete Iscove’s Modified Dulbecco’s Glutamax medium (IMDM; Gibco, 12440061) supplemented with 10% FBS, 50*µ*M β-mercaptoethanol, 5ng/ml IL3 (Peprotech, 213-13). PUER cells were differentiated into macrophages by adding 200nM 4-hydroxy-tamoxifen (OHT; Sigma, H7904-5MG). Cells were differentiated into neutrophils by replacing IL3 with 10ng/ml Granulocyte Colony Stimulating Factor (GCSF; Peprotech, 300-23) and inducing with 100nM OHT after 48 hours. *Gata1*-deficient megakaryocyte-erytrocyte (G1ME) cells were routinely maintained in complete *α*-MEM Glutamax (Gibco, 12561056) supplemented with 20% FBS and 20ng/ml TPO (Peprotech, 315-14).

### Construct design and cloning using Gibson assembly

Putative CRMs were cloned into a pGL4.10*luc2* Luciferase reporter vector (Promega, E6651). The proximal promoter was introduced into the multiple cloning site (MCS) of pGL4.10*luc2* between XhoI and HindIII sites. The distal CRMs were inserted between BamHI and SalI sites downstream of the SV40 late poly(A) signal. CRM sequences are provided in Supplementary File Text S1.

Each CRM or promoter insert was amplified from genomic DNA of C57BL/6J mice using Q5 High-Fidelity 2X Master Mix (NEB, M0492L) following the manufacturer’s instructions. The following PCR cycling conditions were used: initial denaturation of 30s at 98°C, 30 cycles of 30s at 98°C, 30s at 60°C, and 60s at 72°C, and a final extension for 10 minutes at 72°C. Primers included 40bp of sequence homologous to pGL4.10*luc2* (Table S3). Gibson Assembly (GA) reactions (Gibson *et al*., 2009) were carried out using 0.06pmol of digested vector and 0.18pmol of insert, for 60 minutes at 50°C. NEB high-efficiency competent cells (NEB, E5510S) were transformed according to manufacturer’s instructions.

### Transfection and Luciferase assays

PUER or G1ME cells were transfected with a reporter vector and Renilla control vector (pRL-TK, TK promoter, gift of A. Dhasarathy) in a 1:200 ratio using a 4D-Nucleofector (Lonza). PUER cells were transfected with 2.26*µ*g total plasmid DNA in SF buffer (Lonza, V4SC-2096), using program CM134 and incubated for 24 hours prior to luminescence measurement. G1ME cells were transfected with 4.52*µ*g total plasmid DNA in P3 buffer (Lonza, V4SP-3096), using program CM134 and incubated for 6 hours before luminescence measurement. After incubation, Firefly and Renilla luminescence were measured using the Dual-Glo luciferase activity kit (Promega, E2920) and the DTX 880 Multimode Detector (Beckman Coulter) according to manufacturer’s instructions. Transfections were performed in at least 10 replicates.

### Normalization of Firefly luminescence against Renilla luminescence

Well-to-well transfection efficiency variation was controlled for by normalizing Firefly luminescence against Renilla luminescence. Robust errors-in-variables (EIV) regression, implemented according to the method of Zamar (1989), was used to estimate the slope, *β*, of the line *y* = *βx*, where *y* is Firefly luminescence and *x* is the Renilla luminescence. The details of our implementation of robust EIV regression will be described elsewhere (Repele *et al*., 2018). Briefly, *β* was estimated by minimizing the loss function

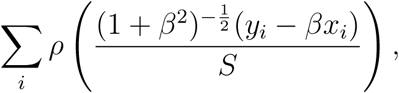

where, *x*_*i*_ and *y*_*i*_ are individual replicates of Renilla and Firefly luminescence measurements, 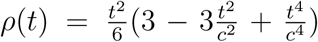 is Tukey’s loss function with *c* = 4.7, and *S* is an estimate of the scale of the residuals. The argument of Tukey’s function is the orthogonal distance of the point (*x*_*i*_, *y*_*i*_) from the regression line. Tukey’s function is bounded for large values of *t*, which limits the contribution of outliers to the loss function and ensures that the slope estimate is robust to outliers. The value of *S* was estimated by solving the equation

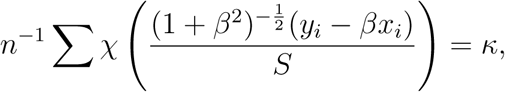

Where *k*= 0.05 and *χ* (*t*) is Tukey’s loss function with *c* = 1.56. The minimization problems were solved by the sequential least-squares quadratic programming (SLSQP) algorithm of the `NLOPTR` package of R, with parameters `xtol_rel` and `maxeval` set to 10^-7^ and 1000 respectively.

95% confidence intervals were estimated by bootstrapping using the R package `BOOT. 999` replicates were subsampled using the ordinary simulation and the function `boot.ci` was used determine confidence intervals using the basic bootstrap method.

### Design and synthesis of mutant CRMs

Mutations to predicted TF binding sites were designed *in silico* with the aid of our sequence-based model of transcription (Bertolino *et al*., 2016). A mutant binding site was created by changing each nucleotide in the wildtype site to one having the lowest frequency in the alignment matrix (Hertz & Stormo, 1999) of the cognate TF (Table S1). The mutated CRM was then simulated in the model to predict its activity and confirm that the targeted site was lost, no new sites had been created, and the other sites were unmodified. If the mutant sequence interfered with other sites or introduced new ones, then nucleotides having the second lowest frequency in the alignment matrix were chosen at a few positions to circumvent interference. Mutant sequences were synthesized using Gibson assembly either with primers carrying the desired mutations or with synthetic dsDNA, or both (Tables S2 and S3). The mutant CRMs were cloned into pGL4.10*luc2* using Gibson assembly as described above. Mutant CRM sequences are provided in Supplementary File Text S1.

### Reverse transcription real-time PCR

Total RNA was extracted using MagJet RNA kit (Thermo, K2731), and reverse transcribed using the High Capacity cDNA Transcription kit (Applied Biosystems, 4368814) following the manufacturer’s instructions. Real-Time PCR was performed using the Ssofast Evagreen Supermix (BioRad, 1725201) in a C1000 Thermal Cycler with CFX384 Real-Time System (BioRad) using the following cycling conditions: initial denaturation of 30s at 95°C followed by 40 cycles of 5s at 95°C and 5s at 60°C. *Cebpa* expression relative to *Hprt* was computed as 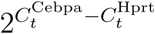, where 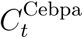 and 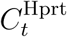 are the threshold cycles for *Cebpa* and *Hprt* respectively. The following primers were used:

*Cebpa*_fwd: ACTTTCCGCGGAGCTGAG

*Cebpa*_rev: ATTTTTGCTCCCCCTACTCG

*Hprt*_fwd: ACCTCTCGAAGTGTTGGATA

*Hprt*_rev: CAACAACAAACTTGTCTGGA

## Results

### The expression of *Cebpa* during macrophage-neutrophil differentiation

We characterized the time course of *Cebpa* expression during the differentiation of PUER cells into macrophages and neutrophils. PUER cells are IL3-dependent hematopoietic progenitors derived from *Spi1*^*-/-*^mice and carry a transgene encoding a PU.1-Estrogen receptor fusion protein (Walsh *et al*., 2002). Uninduced PUER cells function like myeloid progenitors and can be induced to differentiate by treatment with 4-hydroxy-tamoxifen (OHT) into either macrophages or neutrophils in the presence of IL3 or GCSF respectively (Fig. 1B; Dahl *et al*., 2003). For neutrophil differentiation, IL3 medium is completely replaced with GCSF medium 48 hours prior to differentiation.

**Figure 1.**
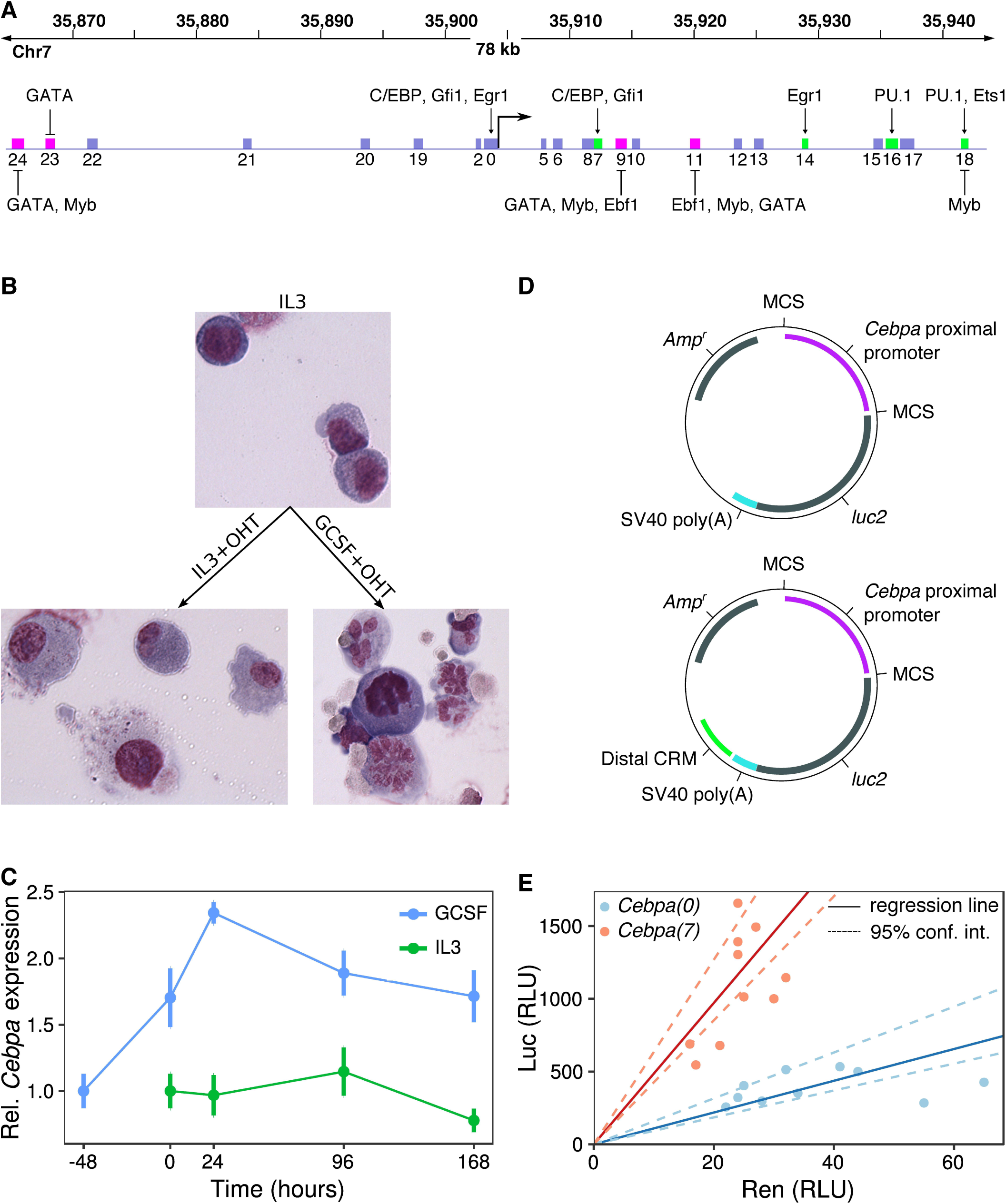
The regulation of *Cebpa* in PUER cells. **A**. A 78kb region surrounding the *Cebpa* TSS is shown. The boxes represent putative CRMs previously identified using evolutionary conservation and analyzed using sequence-based thermodynamic modeling (Bertolino *et al*., 2016). Green and magenta boxes represent enhancers and silencers respectively. The activators and repressors inferred by the model are indicated. **B**. Wright Giemsa stains of PUER cells in uninduced IL3 (top), 7-day OHT-induced IL3 (bottom left), and 7-day OHT-induced GCSF (bottom right) conditions. Uninduced cells have a blast morphology with high nucleocytoplasmic ratio. Cells induced in IL3 conditions have a vacuolated cytoplasm and low nucleocytoplasmic ratio, while induction in GCSF results in cells with segmented nuclei. **C**. Time series of the ratio of *Cebpa* and *Hprt* expression measured by RT-RTPCR during the differentiation of PUER cells. Relative expression has been normalized to average relative expression in uninduced PUER cells. −48 hours and 0 hour points are both measurements from uninduced cells. With the exception of 96 hours GCSF+OHT, for which *N* = 2, *N* ≥3. Error bars show standard error. **D**. Schematics of reporter vectors, based on the pGL4 backbone (Promega), which contain the *Cebpa* promoter immediately upstream of *luc2* either with (below) or without (above) a distal CRM located downstream of the SV40 Poly(A) signal. **E**. Normalization of Firefly luminescence against Renilla luminescence to correct for sample-to-sample variation in transfection efficiency. Points are independent Firefly and Renilla luminescence measurements for *Cebpa(0)* (blue) and *Cebpa(7)* (red). Luminescence is reported in relative luminescence units (RLUs). The ratio of Firefly and Renilla luminescence was estimated as the slope of the best-fit line (solid) determined by robust errors-in-variable (EIV) regression. Dashed lines represent the 95% confidence interval for slope determined by bootstrapping (see Methods).

We measured *Cebpa* gene expression relative to *Hprt* or *Gapdh* using RT-RTPCR in uninduced PUER cells and at four time points during a 7-day course of differentiation in IL3 and GCSF conditions (Fig. 1C). Here we report *Cebpa* expression relative to *Hprt*; similar results were obtained when using *Gapdh* as the reference (data not shown). Overall, *Cebpa* is expressed two-fold higher in GCSF than in IL3 conditions (Wilcoxon rank sum test after pooling time points, N=13, *p* = 2.99 × 10^-6^). *Cebpa* expression increases 70% during the 48 hour pretreatment with GCSF, and another 40% after the first 24 hours of GCSF+OHT treatment. Thereafter, the expression level declines gradually over time to revert to pre-OHT levels at day 7. In contrast, the expression level remains relatively constant after OHT treatment in IL3 conditions. The increased expression of *Cebpa* during neutrophil differentiation is consistent with the essential role that C/EBP*α* plays in neutrophil development and previous analyses of PUER differentiation (Dahl *et al*., 2003; Laslo *et al*., 2006). These data also indicate that most of the regulatory modulation of *Cebpa* expression occurs during the first 24 hours of differentiation.

### The activity pattern of *Cebpa* enhancers

We had previously identified four enhancers of *Cebpa* in a screen utilizing evolutionary conservation and reporter assays (Bertolino *et al*., 2016, Fig. 1A). Three of four enhancers were novel, while one enhancer, *Cebpa(18)*, overlapped with a known enhancer located 37kb downstream of the *Cebpa* TSS (Guo *et al*., 2012, 2014; Cooper *et al*., 2015). Prior to dissecting the *cis*-regulatory logic of these newly identified enhancers, we validated their activity in PUER cells using a statistically robust procedure for measuring reporter activity that we have developed recently (Repele *et al*., 2018).

In transient reporter assays, transfection efficiency can vary over an order of magnitude from sample to sample (Jacobs & Dinman, 2004). In our reporter data, we observed a 2-4 fold variation in luminescence from sample-to-sample (Fig. 1E). The prevalent method of correcting for transfection efficiency variation is to co-transfect an independent reporter, such as the *Renilla* Luciferase expressed from a constitutive promoter, along with the CRM reporter being assayed. Firefly luminescence is then normalized to *Renilla* luminescence to control for sample-to-sample variation in transfection efficiency. Normalizing by taking the ratio of Firefly and *Renilla* luminescence is statistically unsound since it weights low-and high-luminescence replicates equally even though the latter produce more reliable estimates of normalized reporter activity.

In our method, we utilize linear regression to determine the normalized activity as the slope of the best fit line (Fig. 1E), and hence avoid weighting all points equally. Ordinary least squares regression assumes that the values of the independent variable, *Renilla* luminescence in our case, are known exactly and don’t include random errors. Since *Renilla* luminescence is itself a random variable in transient assays, we use robust errors-in-variables (EIV) regression (Casella & Berger, 2001; Zamar, 1989) instead. The estimation of the slope and intercept is rendered insensitive to outliers by utilizing a bounded loss function (Zamar, 1989). Furthermore, the loss function is a sum of the squares of the scaled orthogonal distance of each data point from the line, and hence leads to the minimization of errors in both variables, instead of just the dependent variable. Finally, we performed reporter assays in 10 replicates in order to boost statistical power.

We tested the four previously identified enhancers (Bertolino *et al*., 2016) using this statistically robust methodology. In all reporter data presented in this manuscript, the reporter vectors either carry the *Cebpa* proximal promoter alone (*Cebpa(0)*) or in combination with one of the distal CRMs (Fig. 1D). We denote the vector carrying a CRM along with the promoter as *Cebpa(X)*, where *X* is the CRM number. The comparison of the CRM-bearing reporter with *Cebpa(0)* allows us to discriminate enhancing or silencing CRMs from neutral ones.

All four enhancers upregulated the activity of the promoter robustly and also exhibited cell-type specific patterns of activity (Fig. 2). *Cebpa(7)* is the strongest enhancer in uninduced conditions, upregulating activity ∼6-fold relative to *Cebpa(0)*. *Cebpa(7)*’s enhancing effect is moderated somewhat to 3-fold and 4.5-fold in induced IL3 and GCSF conditions respectively. *Cebpa(14)* has a qualitatively similar activity pattern as *Cebpa(7)*, providing the greatest activation, ∼2.5-fold, in uninduced conditions. *Cebpa(16)* and *Cebpa(18)*, in contrast, have the greatest activity in induced conditions. *Cebpa(16)* upregulates the proximal promoter 4.1-fold in induced GCSF conditions compared to ∼2-fold in uninduced conditions. Similarly, *Cebpa(18)* provides the greatest upregulation of ∼2.5-fold in induced GCSF conditions. Greater activity in induced conditions, when the PU.1-Estrogen receptor fusion protein localizes to nuclei, is consistent with a prediction arising from our previous analysis (Bertolino *et al*., 2016) that *Cebpa(16)* and *Cebpa(18)* are activated by PU.1.

**Figure 2.**
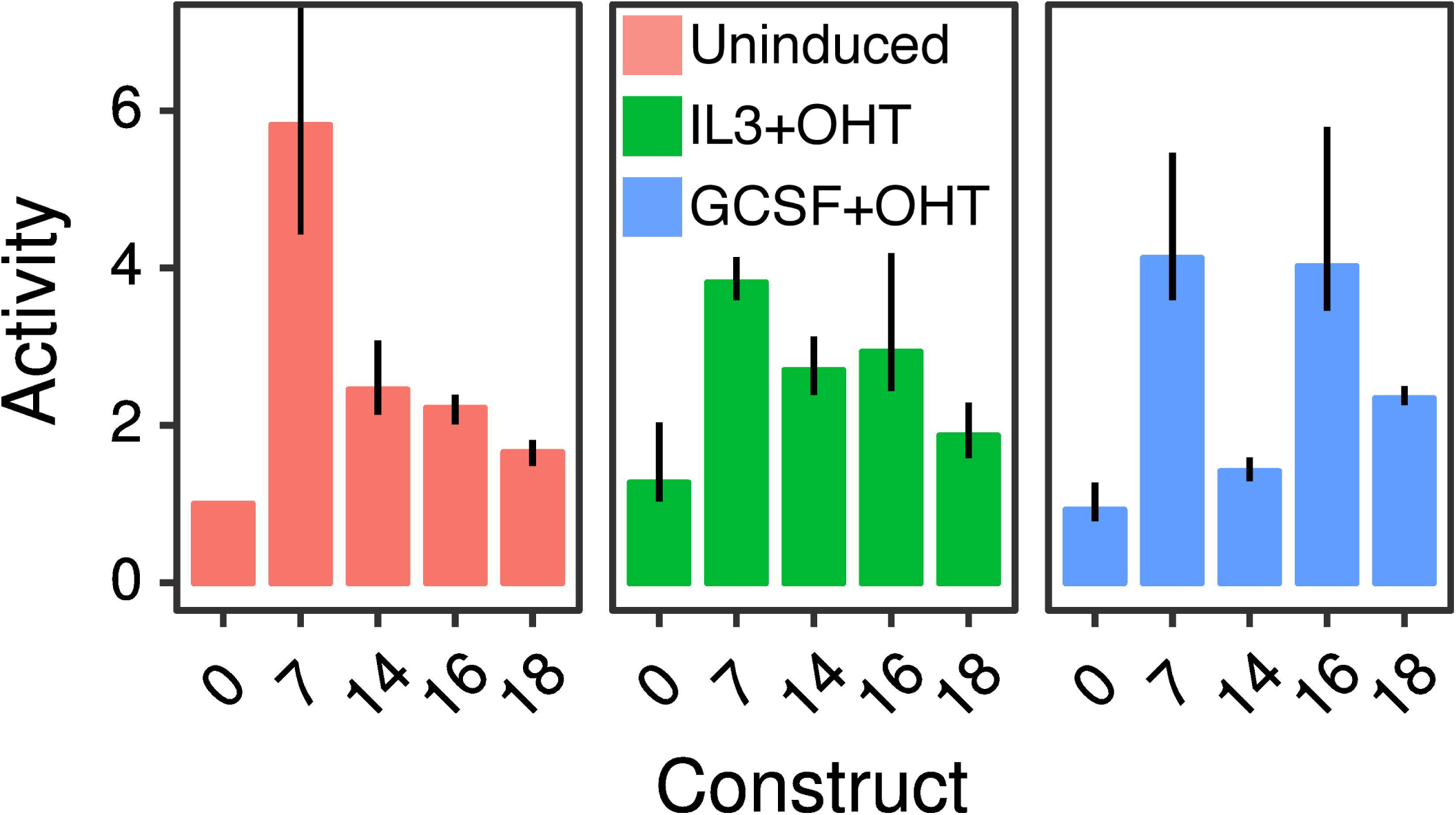
Relative activity of *Cebpa* enhancers in PUER cells. *Cebpa(0)* is the construct bearing the *Cebpa* proximal promoter alone, while the others carry the indicated distal CRM in addition to the proximal promoter. Bar plots show the ratio of construct activity in each condition to *Cebpa(0)* activity in uninduced conditions. Each CRM’s activity was assayed in uninduced (red), induced IL3 (green), induced GCSF (blue) conditions. Reporter assays were performed in 10 replicates.Error bars are 95% confidence intervals. The error bar for *Cebpa(7)* extends to 15.4. Regression plots corresponding to each bar are shown in Figure S1.

### The *cis*-regulatory logic of *Cebpa* enhancers at binding-site resolution

Having rigorously validated the novel enhancers, we next decoded their *cis*-regulatory logic by mutating binding sites predicted by sequence-based models of gene regulation (Bertolino *et al*., 2016). We provide a brief description of the model here and refer the reader to Bertolino *et al.* (2016) for implementation details and equations. Given the DNA sequence of a CRM, the TFs regulating the CRM, and estimates of TF concentrations in one or more conditions, our model predicts the rate of transcription in each condition. The model utilizes position weight matrices (PWMs) to identify binding sites and to compute their binding affinity relative to the consensus site (Berg & von Hippel, 1987). The model then determines the fractional occupancy of each site “thermodynamically” (Segal *et al*., 2008; Kim *et al*., 2013), that is, by enumerating all possible configurations in which the identified sites may be bound. The occupancy of a site in a given configuration takes into account potential cooperative and competitive interactions between TFs. The fractional occupancy is determined by computing a weighted sum over all the configurations in which a particular site is occupied. The model implements position dependent repression, or quenching (Arnosti *et al*., 1996a; Ogbourne & Antalis, 1998; Stopka *et al*., 2005), by reducing the site occupancy of activators bound in a ∼150bp neighborhood of repressor sites. The total strength of a CRM’s interaction with the polymerase holoenzyme complex is determined by computing a weighted sum of individual activator sites’ occupancies, using activation efficiencies as weights. In the penultimate step, crucial for correctly modeling silencers, the model allows for repression over long distances by reducing the interaction strength as a function of repressor site occupancy. In the last step, transcription initiation is modeled with the Arrhenius equation, in which the interaction strength lowers the activation energy barrier, so that greater interaction strength results in higher transcription rates. In summary, the model utilizes well-known biophysical principles and phenomenological rules to predict CRM activity from DNA sequence.

Besides predicting the activity from sequence when the regulating TFs are known, this modeling framework can also be used to learn which TFs regulate a particular CRM and whether they act as activators or repressors. This is achieved by constructing an ensemble of models realizing all possible combinations of the regulatory roles of a set of candidate TFs and identifying which model realization best fits the empirical reporter activity data (Bertolino *et al*., 2016). Whether a particular TF is predicted to act as an activator or repressor is implicit in the combination of regulatory roles represented in the best fitting model. The TFs predicted to regulate each CRM, as well as their binding sites, can be inferred by analyzing the utilization of TFs in the occupancy and interaction-strength calculations of the best fitting model.

Using this reverse engineering methodology, we had inferred a comprehensive map of *Cebpa* CRM regulation at binding-site resolution (Fig. 1A). These inferences, implicit in the internal composition of the best-fit model for each CRM, constitute a set of hypotheses about the *cis*-regulatory logic of *Cebpa*. In order to place the decoded logic on a firm empirical footing, we sought to test these hypotheses by site-directed mutagenesis. The interpretation of site-directed mutagenesis experiments can be challenging because deletions change binding-site spacing while substitutions have the potential to introduce new binding sites. Having CRM models capable of predicting transcription rate from DNA sequence allowed us to circumvent these limitations. For each binding site to be tested, we designed substitutions to abolish binding by choosing the nucleotide least favored at each position according to the PWM of the cognate TF (Hertz & Stormo, 1999). The mutated sequences were simulated in the model to predict their activity. The simulations allowed us to ensure that the mutations did not create any new binding sites for the TFs represented in the model. We tested the decoded logic by synthesizing the mutant CRMs (see Methods), assaying their activity in uninduced and induced PUER cells, and comparing with the theoretical prediction. In what follows, we describe the inferred *cis*-regulatory logic, the predicted effect of mutations, and the empirical results for each enhancer.

#### Cebpa(7)

In the best-fit model, the upregulation of *Cebpa(7)* over *Cebpa(0)* results from activation provided by C/EBP family TFs and Gfi1, which bind 2 and 3 sites respectively (Fig. 3A,B). We tested the predicted sites of the C/EBP family TFs first since C/EBP TFs are known to regulate the proximal promoter (Legraverend *et al*., 1993) and the +37kb enhancer (Cooper *et al*., 2015). We designed a mutant CRM, *Cebpa(7m1)*, which lacks C/EBP sites and is predicted to have half the activity of *Cebpa(7)* in uninduced conditions when simulated in our model (Fig. 3B,C). Next, we synthesized *Cebpa(7m1)* and assayed its activity in both uninduced and induced conditions in PUER cells. We observed a ∼40% reduction of activity in uninduced conditions, matching the model’s prediction and confirming the activation of *Cebpa(7)* by C/EBP family TFs (Fig. 3C,D). The activity was also reduced in induced IL3 conditions, although not to the same extent as was predicted by the model.

**Figure 3.**
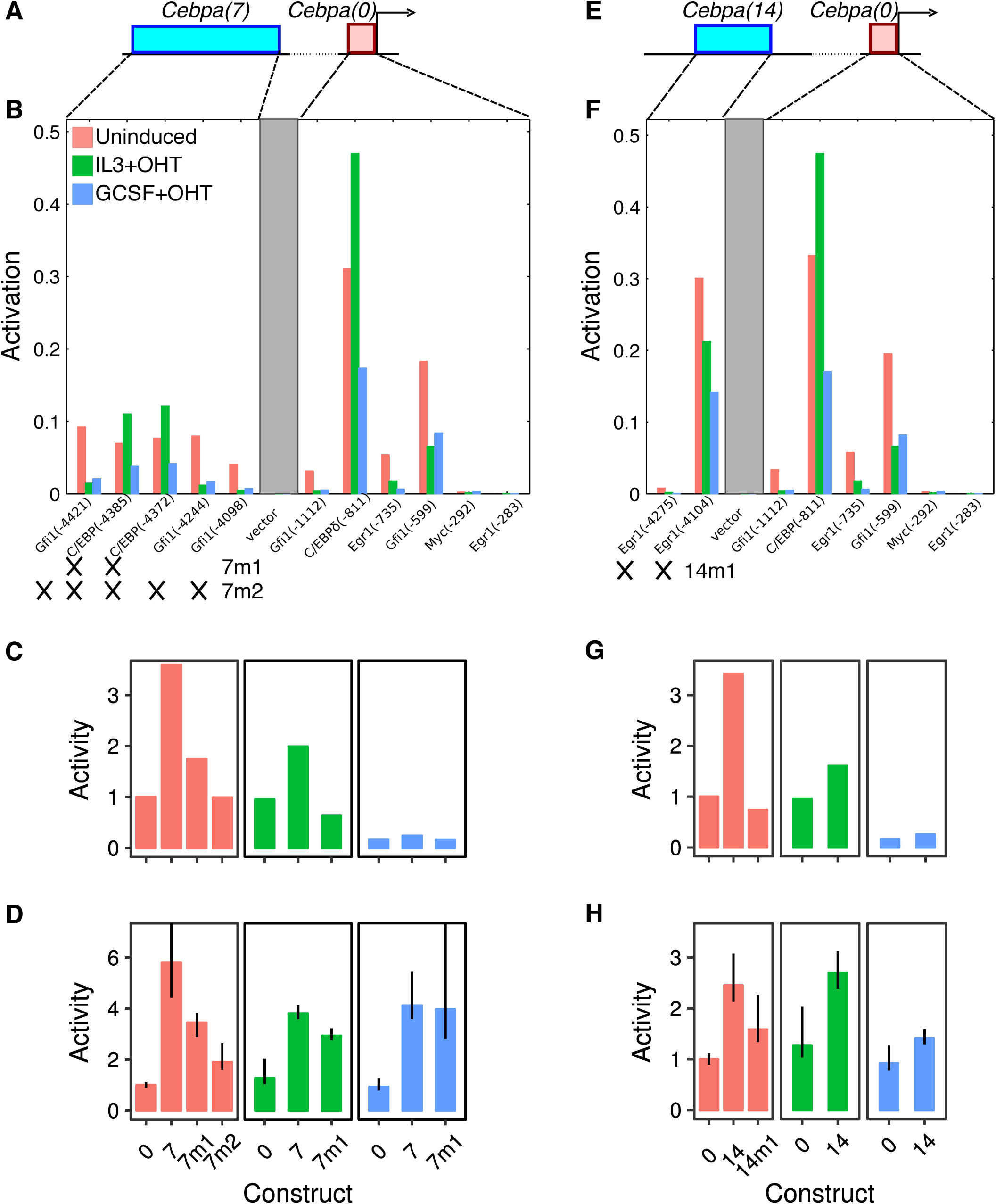
The regulatory logic of *Cebpa(7)* and *Cebpa(14)*. **A–D**. *Cebpa(7)*. **E–H**. *Cebpa(14)*. **A, E**. Schematics of the construct design showing a distal CRM (blue) and the *Cebpa* proximal promoter (red). **B, F**. Activity of each TF activator site predicted by the sequence-based model for each construct. The activity is the amount by which an individual site reduces the activation energy barrier (Bertolino *et al*., 2016) and depends on the occupancy of the site and the efficiency of the bound activator. Sites occurring in the CRM and proximal promoter are shown. The gray box is intervening vector sequence. The *x*-axis shows each binding site modeled and the position of its 5′ end in the reporter construct relative to the 3′ end of the proximal promoter in parentheses.*7m1, 7m2* (panel B), and *14m1* (panel F) refer to mutant constructs tested experimentally. Crosses indicate the sites mutated in each construct. **C, G**. Wildtype and mutant CRM activity predicted by the model *in silico*. **D, H**. Experimentally measured activity of wildtype and synthesized mutant CRMs. Activity levels have been normalized to the activity of *Cebpa(0)* in uninduced conditions. Reporter assays were performed in 10 replicates. Error bars are 95% confidence intervals. Regression plots corresponding to each bar are shown in Figure S2.

The *Cebpa(7m1)* data also suggested that Gfi1 or other as yet unidentified sites are functional since the C/EBP sites did not account for the entirety of *Cebpa(7)* activity. We tested the contribution of Gfi1 sites to the residual activity of *Cebpa(7m1)* by designing a second mutant, *Cebpa(7m2)*, lacking all Gfi1 and C/EBP binding sites (Fig. 3B). Simulations predicted that *Cebpa(7m2)* completely lacked activity in the uninduced condition (Fig. 3C). We observed a further ∼45% reduction in activity compared to *Cebpa(7m1)*, so that *Cebpa(7m2)*’s activity was three-fold lower than that of the wildtype CRM (Fig. 3D). This result confirms the *cis*-regulatory scheme of C/EBP and Gfi1 activation inferred by the model, although residual upregulation of *Cebpa(7m2)* suggests that as yet unknown TFs also contribute to the activity of *Cebpa(7)*.

#### Cebpa(14)

We had inferred that *Cebpa(14)* is activated exclusively by Egr1, which binds the CRM at two predicted sites (Fig. 3E,F). Consistent with regulation by a single factor, Egr1, the activity pattern of *Cebpa(14)* (Fig. 2) matches that of Egr1 (Bertolino *et al*., 2016), having the lowest expression in induced GCSF conditions. We designed a mutant CRM, *Cebpa(14m1)*, which lacks both Egr1 sites. Simulation of *Cebpa(14m1)* predicted a reversion of activity to the level of the proximal promoter (Fig. 3G). Experimentally, we observed a reduction of ∼35% (Fig. 3H), demonstrating the functionality of the Egr1 sites and suggesting that other TFs not represented in the model might also activate *Cebpa(14)*.

#### Cebpa(16)

The model for enhancer *Cebpa(16)* utilizes 4 activator binding sites, 3 for PU.1 and 1 for C/EBP family TFs (Fig. 4A,B). Activation by PU.1 is consistent with the preferential upregulation of *Cebpa(16)* in induced conditions (Fig. 2), when the PU.1-estrogen receptor fusion protein is expected to be localized to the nuclei. A mutant enhancer lacking the PU.1 sites, *Cebpa(16m1)*, was predicted to lack enhancing activity in induced conditions, while being expressed at the same level as wildtype in uninduced conditions (Fig. 4C). Experimentally, *Cebpa(16m1)* behaved as predicted, with an activity nearly half of *Cebpa(16)* and indistinguishable from that of the proximal promoter in induced IL3 conditions (Fig. 4D). There was a much smaller reduction in uninduced conditions so that the activity of *Cebpa(16m1)* was statistically indistinguishable from that of the wildtype enhancer. The activity of *Cebpa(16m1)* was also ∼43% lower than that of *Cebpa(16)* in the induced GCSF condition, although residual upregulation relative to the proximal promoter likely implies that the C/EBP site is also functional.

**Figure 4.**
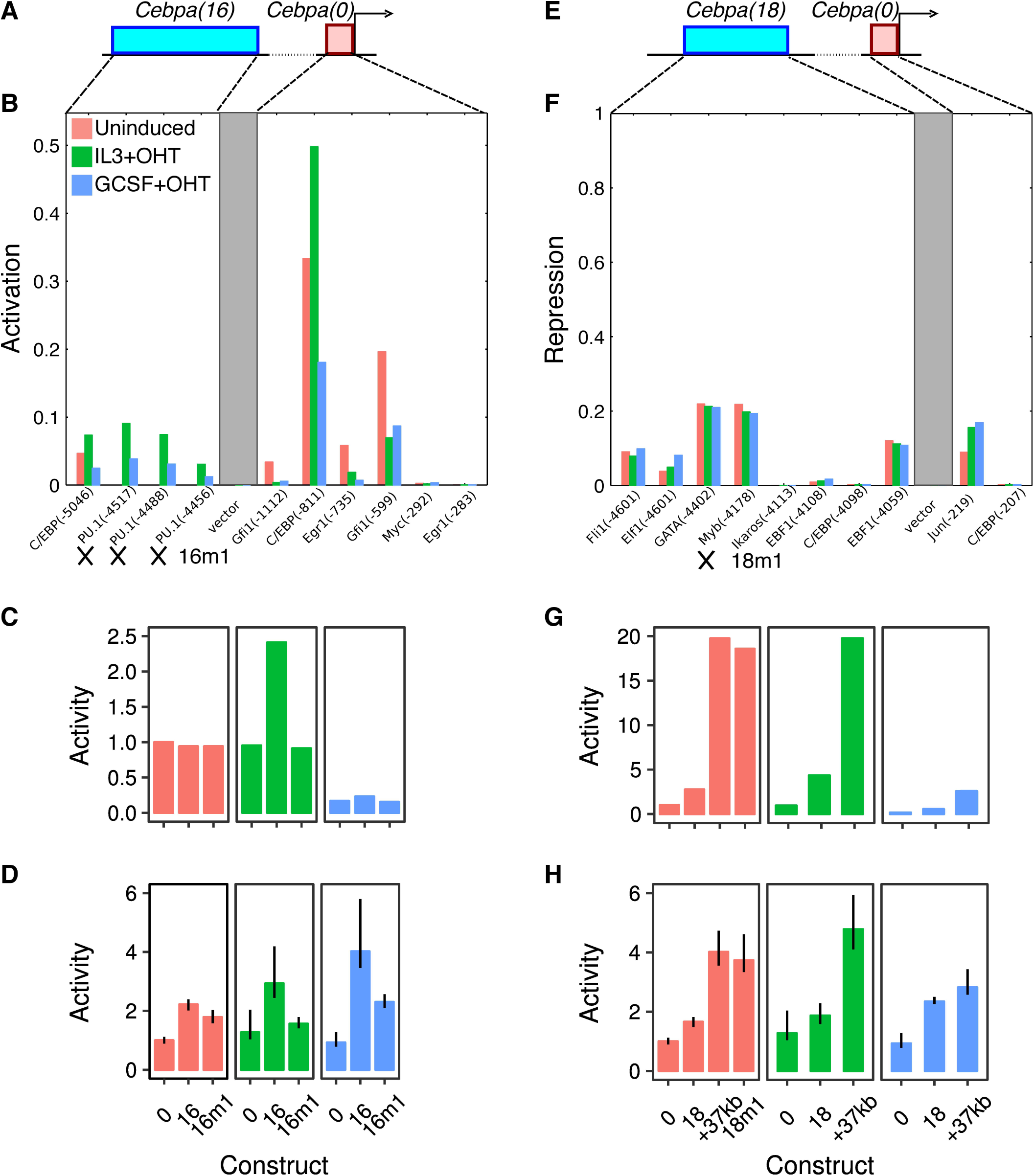
The regulatory logic of *Cebpa(16)* and *Cebpa(18)*. **A–D**. *Cebpa(16)*. **E–H**. *Cebpa(18)*. **A, E**. Schematics of the construct design showing a distal CRM (blue) and the *Cebpa* proximal promoter (red). **B**. Activity of each TF activator site predicted by the sequence-based model for *Cebpa(16)*. *16m1* refers to the mutant construct for testing PU.1 sites (crosses). See the legend of Figure 3B,F for details of the calculations, axes, and legend. **F**. Activity of each TF repressor site predicted by the sequence-based model for *Cebpa(18)*. The repressive activity is the fraction by which the repressor reduces the interaction strength, which results in a higher activation energy barrier. The repressive activity depends on the occupancy of the repressor site and the efficiency of long-range repression of the bound repressor (Bertolino *et al*., 2016). *18m1* refers to the mutant construct for testing the Myb site (cross). See the legend of Figure 3B,F for details of the axes and legend. **C, G**. Wildtype and mutant CRM activity predicted by the model *in silico*. **D, H** Experimentally measured activity of wildtype and synthesized mutant CRMs. Activity levels have been normalized to the activity of *Cebpa(0)* in uninduced conditions. Reporter assays were performed in 10 replicates. Error bars are 95% confidence intervals. Regression plots for *Cebpa(16m1)* are shown in Figure S2. Regression plots for the +37kb enhancer and *Cebpa(18m1)* are shown in Figure S4.

#### Cebpa(18)

We had inferred that *Cebpa(18)* is a PU.1-responsive enhancer with additional activator binding sites for Ets1, Myc, and Gfi1 (Fig. S3C). *Cebpa(18)* (chr7: 35,156,509–35,157,149) encompasses the +37kb *Cebpa* enhancer previously identified by Guo *et al.* (2012) (chr7: 35,156,536– 35,156,974). Site-directed mutagenesis experiments against Ets/PU.1 sites conducted by Cooper *et al.* (2015) independently validated these model predictions in a different cell line, 32Dcl3 myeloid cells.

Given that *Cebpa(18)* is 201bp longer than the +37kb enhancer, we next investigated whether the extra sequences had any function or not. As a first step, we simulated the +37kb enhancer in our model. The model predicted that the activity of the +37kb enhancer is 7-and 4.5-fold higher than that of *Cebpa(18)* in the uninduced and induced IL3 conditions respectively (Fig. 4G), suggesting that the extra sequence has a repressive function. We tested the activity of the +37kb enhancer in PUER cells and observed a ∼2.5-fold increase relative to *Cebpa(18)* in both uninduced and induced IL3 conditions (Fig. 4H), confirming a repressive role for the extra sequence.

We analyzed the repressors predicted by the model to pinpoint the TFs and binding sites responsible for moderating *Cebpa(18)*’s activity. The model had inferred five active repressor sites in *Cebpa(18)*, Fli1, Elf1, GATA, Myb, and Ebf1 (Fig. 4E,F). Of these five, only two, Myb and Ebf1, are unique to *Cebpa(18)*, lying in the extra 201bp of sequence. Of the two TFs, Myb is more likely to mediate the repressive effects since Ebf1 is not expressed in myeloid cells (Pongubala *et al*., 2008). To test the function of Myb, we simulated a mutant CRM lacking the Myb site, *Cebpa(18m1)*, with the model. The model predicted that *Cebpa(18m1)* has a much higher level of activity that is indistinguishable from that of the +37kb enhancer (Fig. 4G). This is indeed how the mutant enhancer behaved in experiment—*Cebpa(18m1)*’s activity was derepressed relative to *Cebpa(18)* and indistinguishable from that of the +37kb enhancer (Fig. 4H). Since Myb is down-regulated in induced PUER cells (Bertolino *et al*., 2016), this result suggests that the upregulation of *Cebpa* enhancers in induced conditions is a consequence not just of a gain in activation by PU.1, but also a loss of repression by Myb.

To summarize, we have decoded the *cis*-regulatory logic of four *Cebpa* enhancers at the resolution of individual binding sites. In all cases, the model’s predictions were borne out by experiment. The identified TFs and their sites are likely the most important regulators of *Cebpa* during macrophage-neutrophil differentiation since they account for most of the CRM activity. The investigated TFs do not however account for all of the CRM activity, suggesting that other TFs not represented in the model also perhaps regulate *Cebpa*. The overall picture that emerges is that C/EBP family TFs, Gfi1, and Egr1 support *Cebpa*’s expression in uninduced or progenitor conditions by binding to *Cebpa(7)* and *Cebpa(14)*. Activation in induced conditions is provided via *Cebpa(16)* and *Cebpa(18)* by increased PU.1 activation and a loss of Myb repression.

### The role of novel silencer elements in hematopoietic lineage resolution

Our previous analysis had revealed CRMs that, when placed in the reporter vector along with the *Cebpa* promoter, reduced the activity of the construct to levels lower than that of the promoter alone (Bertolino *et al*., 2016). This mode of action is consistent with the definition of silencers (Ogbourne & Antalis, 1998; Davidson, 2006). The reduction of activity to levels lower than that of the promoter alone implies that the repressors binding to these silencers act in a dominant manner, similar to long-range repression observed in *Drosophila* (Dunipace *et al*., 2011; Perry *et al*., 2011). Furthermore, the CRMs in question, *Cebpa(9), Cebpa(11), Cebpa(23)*, and *Cebpa(24)*, lie 9–40kb away from the *Cebpa* TSS (Fig. 1A), implying that dominant repression occurs over long distances. Our analysis had inferred that the silencers were repressed by GATA family TFs, Ebf1, and Myb (Fig. 1A; Bertolino *et al*., 2016). Gata1/Gata2 and Ebf1 play key roles in the specification of the red-blood cell and Bcell lineages respectively (Cantor & Orkin, 2002; Pongubala *et al*., 2008),while Myb has been implicated in megakaryocyte development (Lu *et al*., 2008). These inferences are supported by evidence that Gata2 binds to *Cebpa(11)* and *Cebpa(24)* in G1ME cells (Bertolino *et al*., 2016, Fig. S11A), which are blocked at the MEP stage and can be differentiated into erythrocytes (Stachura *et al*., 2006; Doré *et al*., 2012).

These observations motivated the hypothesis that was the subject of our subsequent experiments. We hypothesized that dominant repression mediated via distal silencers is a mechanism for resolving hematopoietic lineages. We tested this hypothesis by 1) checking whether the silencers do, in fact, exert dominant repression in a non-myeloid cell type, and 2) determining whether the silencing is attributable to the predicted repressors.

### The activity pattern of *Cebpa* silencers

Before testing the activity of the silencers in a non-myeloid cell type, we measured their activity in PUER cells using the statistically rigorous methodology we developed for analyzing reporter data. In PUER cells, all of the silencers were either neutral or had weak enhancing activity (Fig. S5). This result implies that previous observations of reduced activity in PUER cells (Bertolino *et al*., 2016) were likely artifacts of low sample size or statistically unsound normalization. Furthermore, this result implies that these CRMs do not silence *Cebpa* in a myeloid background.

Next, we tested the function of silencers in G1ME cells, representative of the red-blood cell lineage. We chose the red-blood cell lineage since Gata2 is known to bind the silencers *Cebpa(11)* and *Cebpa(24)* in G1ME cells (Bertolino *et al*., 2016, Fig. S11A). Even though we could not detect *Cebpa* expression in G1ME cells (Fig. S8), the *Cebpa* promoter had detectable activity (Fig. 5). This suggested that additional repression is required to completely silence *Cebpa*. Reporter vectors carrying *Cebpa(9), Cebpa(11), Cebpa(24)* in addition to the promoter had ∼3-fold lower activity compared to the promoter alone (Fig. 5). The silencers, therefore, while being inert in myeloid cells, repress the *Cebpa* proximal promoter in the red-blood cell lineage.

**Figure 5.**
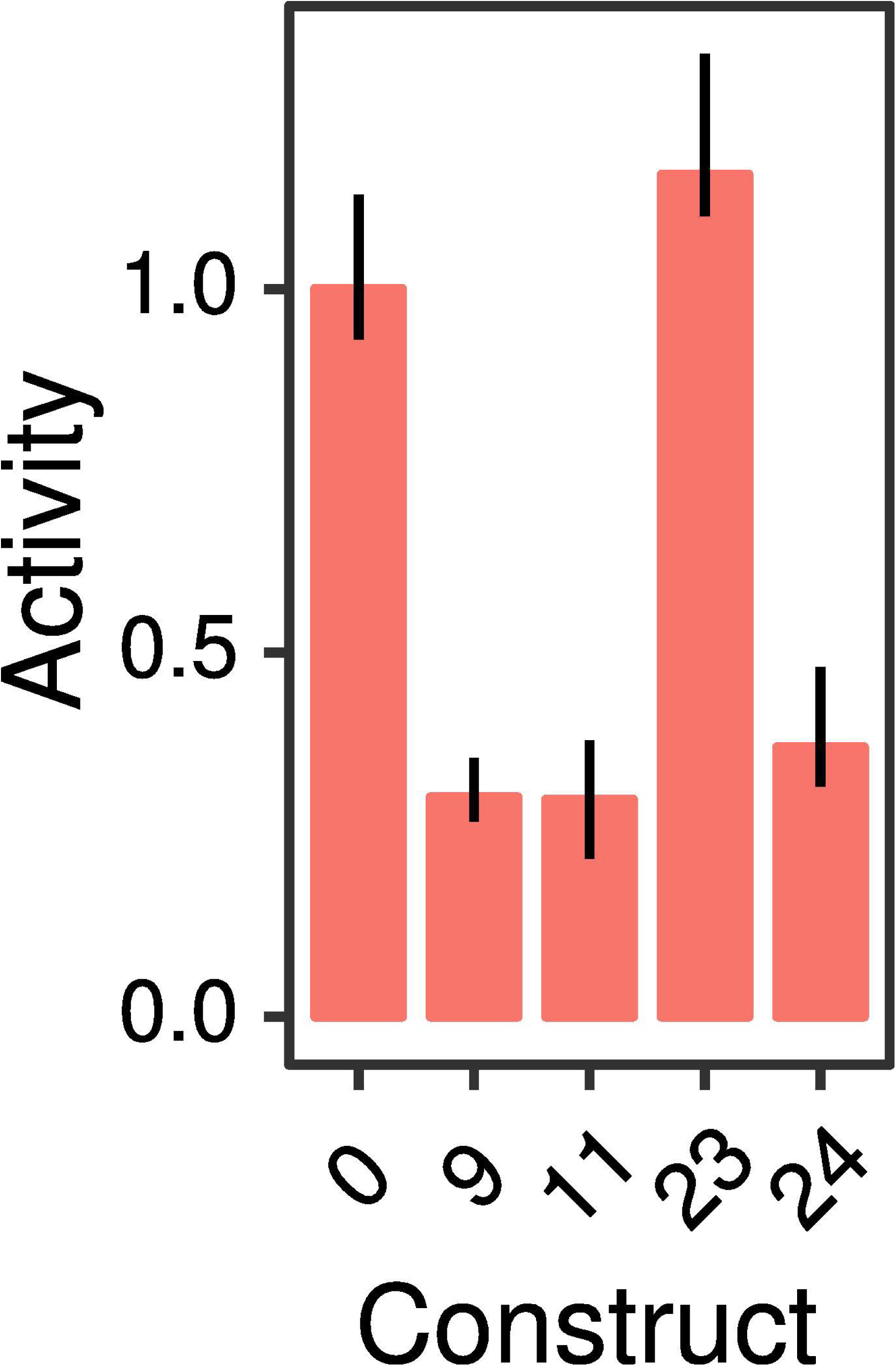
Relative activity of *Cebpa* silencers in G1ME cells. *Cebpa(0)* is the construct bearing the *Cebpa* proximal promoter alone, while the others carry the indicated distal CRM in addition to the proximal promoter. Bar plots show the ratio of each construct’s activity to *Cebpa(0)*. Reporter assays were performed in 10 replicates. Error bars are 95% confidence intervals. Regression plots corresponding to each bar are shown in Figure S7.

### GATA and Myb repress the *Cebpa* proximal promoter in a dominant and redundant fashion

We decoded the *cis*-regulatory logic of the silencers using the same model-guided strategy as was employed for the enhancers. In contrast to the enhancers, each of which had a distinctive regulatory scheme, the validated silencers shared a common regulatory motif. All three silencers had GATA and Myb sites, which were predicted to be among the most active in each silencer (Fig. 6B,E,H). GATA family TFs and Myb are plausible repressors of *Cebpa*. Knocking down Gata2 leads to the derepression of *Cebpa* in G1ME cells (Huang *et al*., 2009), while we have demonstrated that Myb represses *Cebpa(18)* (Fig. 4G,H). We synthesized mutants CRMs—*Cebpa(9m1), Cebpa(11m1)*, and *Cebpa(24m1)*—lacking binding sites for both TFs (Fig. 6B,E,H). *Cebpa(11m1)* carried additional mutations in an Ebf1 site but was functionally equivalent to a GATA/Myb mutant since Ebf1 is not expressed in the red-blood cell lineage (http://biogps.org/gene/13591; Hagman *et al*., 1993; Doré *et al*., 2012). As predicted by the model, the mutant CRMs were derepressed relative to wildtype and, in the case of *Cebpa(9m1)* and *Cebpa(11m1)*, their activity was indistinguishable from that of *Cebpa(0)* (Fig. 6C,F,I). This implies that the silencing can be attributed specifically to GATA and Myb, which account for the entirety of the effect in two of three silencers.

**Figure 6.**
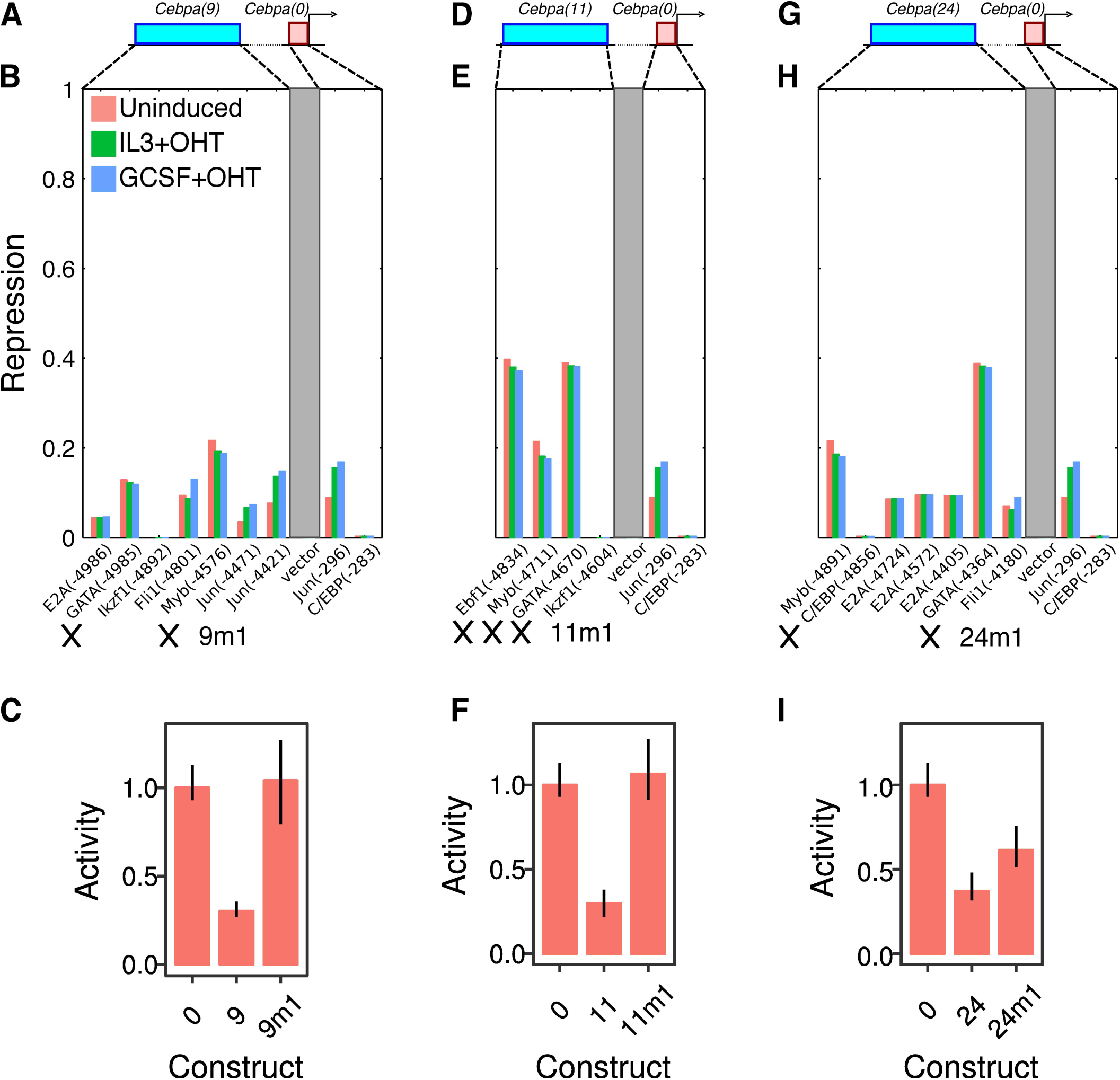
Silencing relies on a GATA/Myb motif shared by functional *Cebpa* silencers. **A–C**. *Cebpa(9)*. **D–F**. *Cebpa(11)*. **G–I**. *Cebpa(24)*. **A, D, G**. Schematics of the construct design show-ing a distal CRM (blue) and the *Cebpa* proximal promoter (red). **B, E, H**. The activity of the TF repressor sites predicted by the model for each silencer. See the legend of Figure 4F for details of the calculations, axes, and legend. *9m1, 11m1*, and *24m1* refer to mutant CRMs and crosses indicate mutated sites. **C, F, I**. Experimentally measured activity of wildtype and synthesized mutant silencers in G1ME cells. Activity levels have been normalized to the activity of *Cebpa(0)*. Reporter assays were performed in 10 replicates. Error bars are 95% confidence intervals. Regression plots corresponding to each bar are shown in Figure S9.

Next, we investigated how GATA and Myb jointly repress the activity of the *Cebpa* proximal promoter. We considered three hypotheses and tested them by mutating GATA and Myb sites individually in *Cebpa(11)* (Fig. 7A). First, it is possible that GATA and Myb repress the proximal promoter redundantly (Delneri *et al*., 1999), so that only one functional site is sufficient to achieve silencing. The second possibility is that of synergism (Arnosti *et al*., 1996b; Heinz *et al*., 2010; Akasaka *et al*., 2001), in which GATA and Myb would have much greater silencing activity together than individually. The third possibility is that of context-dependent role switching, when a TF switches its role when bound near a second TF. For example, in the *Drosophila* blastoderm, the repressor Hunchback activates gene expression when bound near Bicoid (Small *et al*., 1991; Simpson-Brose *et al*., 1994). We tested two new constructs, *Cebpa(11m2)* and *Cebpa(11m3)*, which carry impaired GATA or Myb sites respectively. Both of the constructs carrying only one functional repressor site were able to silence the proximal promoter (Fig. 7C). This result provides strong support for the hypothesis that GATA and Myb are capable of repressing the promoter individually and act redundantly.

**Figure 7.**
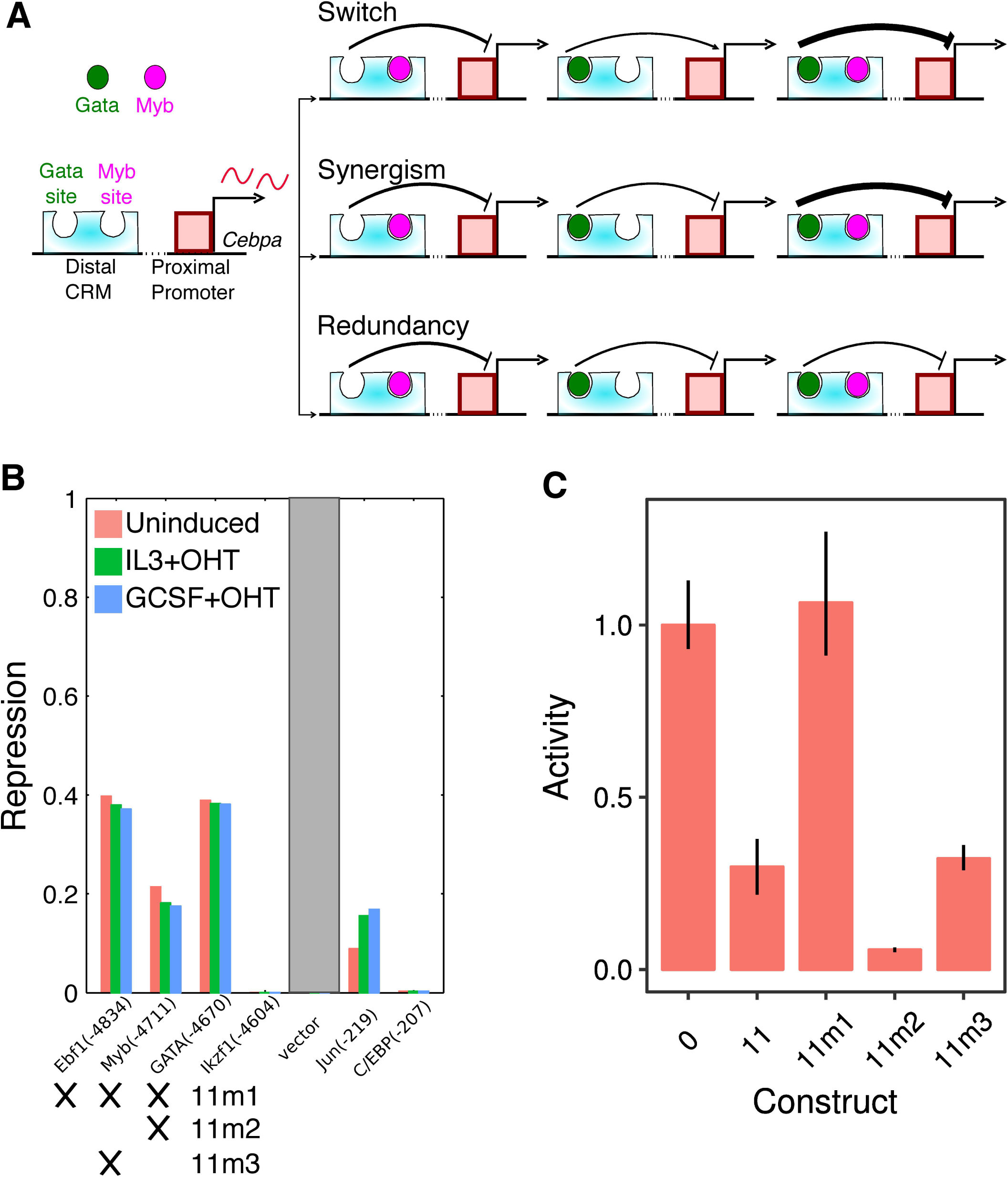
GATA and Myb repress the *Cebpa* proximal promoter redundantly. **A**. Three potential hypotheses for the combined silencing of the promoter by GATA and Myb. **B**. The activity of the TF repressor sites predicted by the model for each silencer. See the legend of Figure 4F for details of the calculations, axes, and legend. *11m1, 11m2*, and *11m3* refer to tested mutant CRMs and crosses indicate mutated sites. **C**. Experimentally measured activity of wildtype and synthesized mutant silencers in G1ME cells. Activity levels have been normalized to the activity of *Cebpa(0)*. Reporter assays were performed in 10 replicates. Error bars are 95% confidence intervals. Regression plots corresponding to each bar are shown in Figure S9.

In summary, we have shown that three of four putative silencers are capable of attenuating the activity of the *Cebpa* proximal promoter in a dominant manner. Dominant repression only occurs in the red-blood cell lineage and the CRMs do not silence the proximal promoter in myeloid cells. Lastly, the silencing activity is attributable to a regulatory motif shared by all three silencers— GATA and Myb sites that act redundantly.

## Discussion

We have comprehensively analyzed the regulation of 7 CRMs neighboring *Cebpa* at the resolution of individual binding sites. In the process of doing so, we have also verified the predictive ability of a thermodynamic model of mammalian gene regulation that we developed recently (Bertolino *et al*., 2016). It is worth noting that prior to our efforts, thermodynamic modeling was limited to *Drosophila* gene regulation (Reinitz *et al*., 2003; Janssens *et al*., 2006; Zinzen *et al*., 2006; Segal *et al*., 2008; Kazemian *et al*., 2010; Kim *et al*., 2013; Samee & Sinha, 2014; Barr & Reinitz, 2017), with a single gene, *even-skipped*, as the focus of most of the work. Our model is closely related to its *Drosophila* counterparts, incorporating just one additional mechanism, long-distance dominant repression, lacking in the latter. The ability of models with shared mechanisms of gene regulation to predict reporter activity in these divergent species supports the view that the rules of transcriptional regulation are universal.

Long-distance dominant repression was required to correctly model silencers (Bertolino *et al*., 2016) and our results here suggest that it is a novel means of resolving hematopoietic lineages. During differentiation, multipotential cells are known to promiscuously and stochastically express genes from multiple daughter lineages, a phenomenon known as multilineage priming (Hu *et al*., 1997; Månsson *et al*., 2007; Velten *et al*., 2017). A natural corollary is that error-correcting mechanisms must exist to repress this ectopic expression once differentiation toward a particular lineage has commenced. Two main error-correcting mechanisms have been found so far. First, a TF may directly inhibit an alternative lineage activator from inducing its targets by protein-protein interaction. For example, during erythroid development, Gata1 prevents PU.1 from activating its targets by binding to PU.1 and displacing cJun, it’s co-activator (Cantor & Orkin, 2001). The second method involves transcriptional repression by a TF that binds to the promoter of a gene expressed in alternative lineages, such as the bistable switch between Egr1/2 and Gfi1 that operates during macrophage-neutrophil development (Laslo *et al*., 2006).

We hypothesized that silencers contribute to lineage resolution since our model had inferred TFs known to promote the development of non-myeloid lineages, GATA (Doré & Crispino, 2011), Myb (Lu *et al*., 2008), and Ebf1 (Pongubala *et al*., 2008), as effectors of silencer-mediated repression (Bertolino *et al*., 2016). This hypothesis predicts that the silencers of *Cebpa* would have much lower activity in the myeloid lineage, where the gene is expressed, than in non-myeloid ones, where it is not expressed. When we assayed the activity of the silencers in PUER cells, which belong to the myeloid lineage, we found they they acted as enhancers or neutral elements (Fig. S5), confirming part of the prediction. In contrast, 3 of 4 putative silencers downregulated promoter activity ∼3-fold in G1ME cells (Fig. 5) belonging to the megakaryocyte-erythrocyte lineage, confirming the rest of the prediction. Furthermore, silencing by distal elements appears to be necessary for quenching *Cebpa* expression in G1ME cells since the promoter has detectable activity (Fig. 5), even though *Cebpa* expression is not detectable in G1ME cells (Fig. S8).

The mode of silencer action, dominant repression over long distances, is conceptually similar in function to long-distance repression mechanisms inferred in other developmental systems. *Drosophila* segmentation genes (Dunipace *et al*., 2011; Perry *et al*., 2011) as well as murine *fgf8* (Marinić *et al*., 2013) are regulated by multiple redundant enhancers, also called shadow enhancers, which are coexpressed in overlapping spatiotemporal domains. In reporter assays, individual elements often exhibit ectopic expression that is corrected when the reporter gene is driven by multiple redundant CRMs together. Based on such experiments, Perry *et al.* (2011) proposed that long-range repressors, such as Tailless or Huckebein, bind to one of the redundant elements to repress activity driven by the other. Ectopic expression was restored when the multiple-CRM transgene was crossed into a genetic background lacking Tailless and Huckebein expression (Perry *et al*., 2011). Our site-directed mutagenesis experiments with *Cebpa*, in which dominant repression is completely abolished upon mutating Gata and Myb sites (Fig. 6), echo this result from *Drosophila*, and suggest that long-range dominant repression is a conserved mechanism for tight control of gene expression programs during development.

Detailed analysis of the regulatory logic of silencers revealed two layers of redundancy in their function. First, structural similarity underlies the functional equivalence of all three silencers. All silencers contain the same regulatory motif, a pair of GATA and Myb sites, that mediates dominant repression of the *Cebpa* promoter (Fig. 6). Second, the regulatory motif itself is structured redundantly since mutating either GATA or Myb alone is not sufficient for relieving dominant silencing (Fig. 7). We did not find evidence for synergy, where the combined effect of the two sites is greater than the sum of individual effects, implying that GATA and Myb function redundantly. The presence of multiple functionally equivalent and structurally homologous silencers in the locus suggests that the regulatory architecture of distal silencing is similar to that of distal activation by multiple redundant enhancers (Hong *et al*., 2008; Frankel *et al*., 2010). Redundant enhancers have been shown to ensure robust and precise gene expression (Perry *et al*., 2010, 2011, 2012), leading us to speculate that redundant silencing might serve a similar function.

The activation of *Cebpa* CRMs in myeloid cells also occurs in a redundant and overlapping regulatory arrangement. All enhancers are simultaneously co-active in nearly all conditions tested in PUER cells. The sole exception is CRM18, for which we could not detect statistically significant upregulation in induced IL3 (macrophage) conditions (Fig. 2). Shadow enhancers were first identified in genomewide ChIP data as distal binding sites of the regulators of classical enhancers (Hong *et al*., 2008). Similarly, the coactive enhancers of *Cebpa* share common regulators. *Cebpa(16)* (Fig. 4B) and *Cebpa(18)* (Fig. S3) are activated by PU.1 and other ETS factors, while CRM7 and CRM16 are activated by C/EBP*α*. CRM14 is the exception with predicted and verified binding sites for Egr1 (Fig. 3F) unique to itself.

The overall picture that emerges from our analysis of *Cebpa* enhancers and silencers is that of a distributed and specialized control scheme (Fig. 1). CRMs distributed over an ∼80kb region specialize in either activation or repression. Specialization is a departure from *cis*-regulatory organization of *Drosophila* segmentation genes, whose enhancers are capable of both activation and repression (Small *et al*., 1993; Gray *et al*., 1994; Perry *et al*., 2011; Dunipace *et al*., 2011). Although this arrangement could be evolutionary happenstance, it is also possible that it serves a functional purpose. Despite the antagonism between *Gata1*/*Gata2* and *Cebpa* that we (Bertolino *et al*., 2016, Fig. 7) and others (Doré *et al*., 2012) have demonstrated, Gata2 and *Cebpa* are known to be co-expressed in eosinophils, basophils, and mast cells (Iwasaki *et al*., 2006). The regulatory logic of *Cebpa*, therefore, must allow for expression even in the presence of Gata2 protein. We propose that separable activation and silencing allows *Cebpa* to be expressed at intermediate levels in eosinophils, basophils, and mast cells. Under this hypothesis, the enhancers are active in all GMP-derived cells, while Gata2-dependent silencers are active in MEPs and the subset of myeloid cells where Gata2 is expressed. The quenching of *Cebpa* expression in the red-blood cell lineage is the combined result of the induction of silencing by Gata2/Myb and a lack of activation. In Gata2 expressing myeloid cells, both enhancers and silencers are simultaneously active, resulting in an intermediate level of expression of *Cebpa*. The hypothesis makes the readily testable prediction that both the enhancers and silencers of *Cebpa* should be active in eosinophils, basophils, and mast cells.

Our work raises several questions about gene regulation during development that likely warrant further investigation. First, is dominant repression mediated through distal silencers a general means of resolving gene expression programs during cell-fate specification? The relative paucity of well-characterized silencers might reflect a lack of attention rather than an intrinsic deficit in their abundance or functional importance. Screens (Arnold *et al*., 2013; Yáñez-Cuna *et al*., 2014) for, as well as detailed analyses of, regulatory elements have largely focused on enhancers so far. Large-scale screens modeled on STARR-seq (Arnold *et al*., 2013) might be able to reveal if silencers are more common than previously thought. Another problem highlighted by the multiplicity of co-expressed enhancers in the *Cebpa* locus is how multiple CRMs jointly regulate a target gene. Classical enhancers (Banerji *et al*., 1981) have long been regarded as acting independently—additively—in a distance independent manner. Recent quantitative experiments with loci having two enhancers, however, do not support the classical assumptions and reveal non-additive behavior (Perry *et al*., 2011; Dunipace *et al*., 2011; Webber *et al*., 2013; Marinić *et al*., 2013; Bothma *et al*., 2015), including sub-and super-additive responses. A strictly modular *cis*-regulatory architecture cannot lead to non-additive responses, explaining which requires that we postulate that CRMs interact or influence each others’ activities. Understanding gene regulation underlying cell-fate decisions will therefore require that we discover how CRMs influence each others’ activities and model loci as interconnected complex systems.

## Acknowledgments

PUER cells were a gift of R. Dahl. G1ME cells were a gift of M. Weiss. We thank R. Dahl, D. Darland, T. Rhen, S. Nechaev, and A. Dhasarathy for discussion and technical assistance and T. Rhen for comments on the manuscript. This work was supported by the National Science Foundation under Grant Nos. 1615916 and IIA-1355466, project UND0019821 (North Dakota EPSCoR).

